# The Neotropical endemic liverwort subfamily Micropterygioideae had circum-Antarctic links to the rest of the Lepidoziaceae during the early Cretaceous

**DOI:** 10.1101/2023.11.16.567484

**Authors:** Antonio L. Rayos, Matthew A. M. Renner, Simon Y. W. Ho

## Abstract

Lepidoziaceae are the third-largest family of liverworts, with about 860 species distributed on all continents. The evolutionary history of this family has not been satisfactorily resolved, with taxa such as Micropterygioideae yet to be included in phylogenetic analyses. We inferred a dated phylogeny of Lepidoziaceae using a data set consisting of 13 genetic markers, sampled from 147 species. Based on our phylogenetic estimate, we used statistical dispersal-vicariance analysis to reconstruct the biogeographic history of the family. We inferred a crown age of 197 Ma (95% credible interval 157–240 Ma) for the family in the Australian region, with most major lineages also originating in the same region. Micropterygioideae are placed as the sister lineage to Lembidioideae, with these two groups diverging from each other about 132 Ma in the South American–Australian region. Our results suggest a circum-Antarctic link between Micropterygioideae and the rest of the family, along with extinction of the lineage in the region. Crown Micropterygioideae were inferred to have arisen 45 million years ago in South America, before the continent separated from Antarctica. Our study reveals the influence of past geological events on the evolution and distribution of a widespread and diverse family of liverworts.

## Introduction

Liverworts (Marchantiophyta), one of the three major groups of bryophytes, emerged soon after the colonization of terrestrial habitats by plants during the Ordovician, but the lineage underwent a marked diversification during the early Paleogene (Simpson, 2010; Vanderpoorten and Goffinet, 2009). Among the most widely distributed families of liverworts are Lepidoziaceae, which are the third-largest liverwort family and comprise about 860 species in 29–31 genera and seven subfamilies (Crandall-Stotler et al., 2009; Cooper et al. 2011; Cooper, 2013). As with other diverse liverwort families, many genera of Lepidoziaceae are understood to have arisen in the early Cenozoic (Cooper et al. 2012). Representatives of Lepidoziaceae occur on all continents and inhabit a wide variety of bioclimatic zones, habitat types, and substrates, including soil, peaty ground, decaying wood, and tree trunks.

While exhibiting an almost unparalleled diversity of form in the gametophyte generation, species of Lepidoziaceae are unified by a set of unique morpho-anatomical characteristics of the sporophyte, including very small spores, elaters with blunt ends, and two-phase ontogeny of the capsule epidermis (Schuster, 1969, 2000). All members of the family also share isophyllous gynoecial branches (Schuster, 2000). Although the monophyly of the family *sensu* Schuster (2000) has been established with confidence, molecular phylogenetic studies have not completely resolved the evolutionary relationships among extant species. The first molecular phylogenetic analysis of Lepidoziaceae used three organellar markers and revealed polyphyly in the subfamilies Zoopsidoideae, Lepidozioideae, and Lembidioideae (Heslewood and Brown, 2007). Subsequently, analysis of a larger data set comprising 10 loci from 93 species confirmed the polyphyly of Lepidozioideae and Zoopsidoideae and recovered Lembidioideae as paraphyletic (Cooper et al., 2011). The paraphyly of Lembidioideae motivated the transfer of *Kurzia* to the subfamily from Lepidozioideae (Cooper, 2013). However, sequence data from 10 molecular markers did not allow confident estimation of the basal relationships among subfamilies of Lepidoziaceae, and these remain unresolved. In addition, several distinct taxa, including the subfamily Micropterygioideae, were not included in the molecular phylogenetic studies of Cooper et al. (2011), and their phylogenetic relationships remain opaque.

Micropterygioideae contain two genera, *Micropterygium* and *Mytilopsis*, which differ from the other subfamilies in their conduplicate leaves that have an abaxial wing or ridge (Schuster, 1969; Schuster, 2000). In some of their features, the subfamily has morphological similarities to some members of Lembidioideae. *Micropterygium* includes about 20 species (with a representative species, *M. leiophyllum*, shown in Figure 1) and differs from the monotypic *Mytilopsis* in having underleaves and lateral-intercalary rather than ventral-intercalary vegetative branches. The Micropterygioideae are confined to the Neotropics, where species mostly occur in the lowlands, although some species reach the Andes (up to 3140 m above sea level). Most species are limited to the Amazonian drainage, but some extend to the Caribbean Islands.

**Fig. 1.**
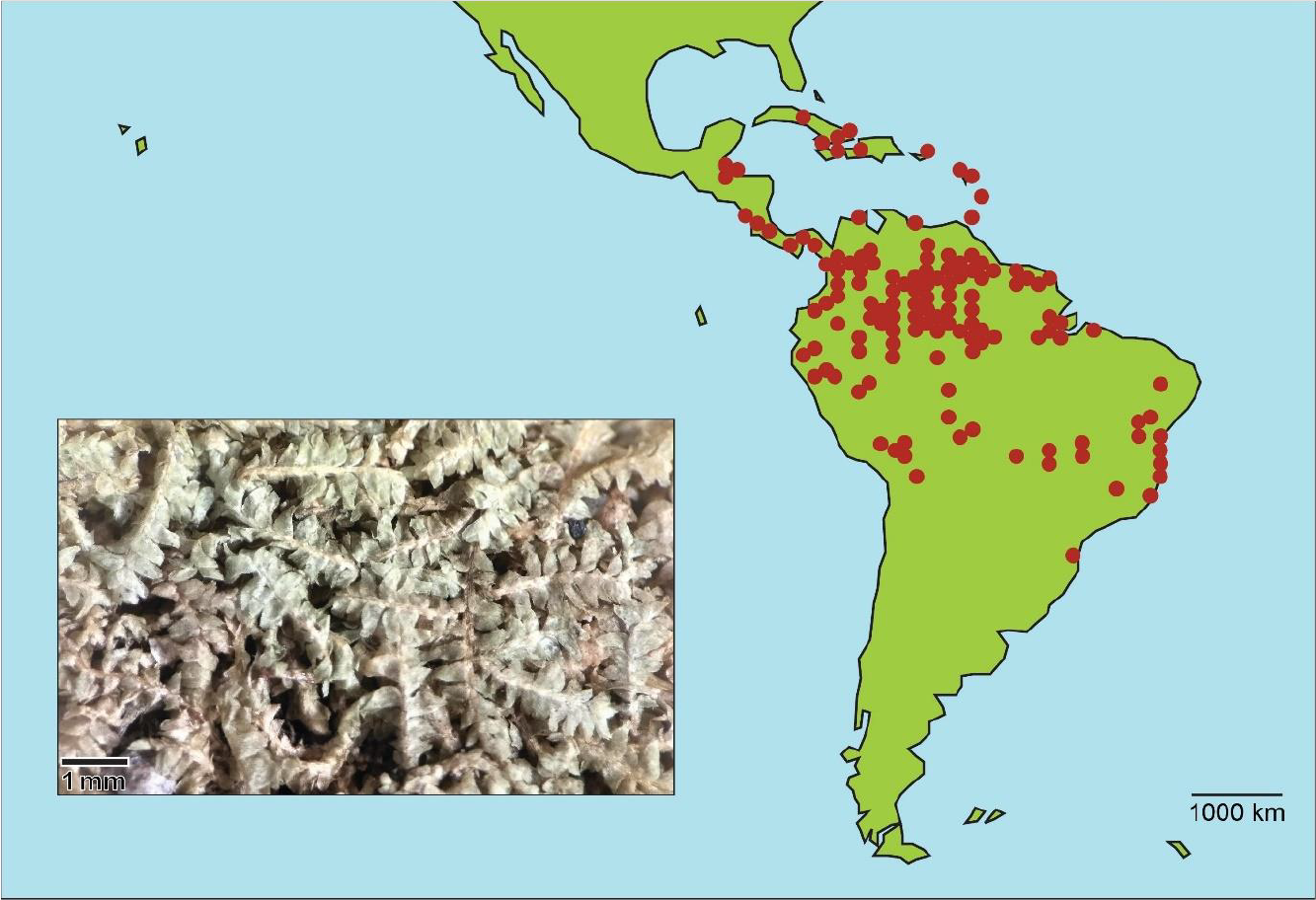
Geographic distribution of Micropterygioideae, with a photomicrograph of a representative species (*Micropterygium leiophyllum*). The red dots represent records in GBIF: *Micropterygium* Lindenb., Nees & Gottsche, and *Mytilopsis albifrons* Spruce (GBIF Backbone Taxonomy, checklist data set https://doi.org/10.15468/39omei).

Lepidoziaceae show generic richness and high endemism concentrated in the circum-Antarctic region, a pattern that has been explained by an origin, or at least a diversification, centred on Gondwana (Schuster, 2000). The confinement of Micropterygioideae to the Neotropics is unusual and could be explained by extinction of the lineage in the circum-Antarctic region. This is not impossible, given that micro- and macrofossils of vascular plants from southern temperate regions suggest that extinction is a general biogeographic feature of all regions, e.g., Winteraceae in South Africa (Coetzee and Praglowski, 1988), *Casuarina* and *Eucalyptus* in New Zealand (Mildenhall, 1980), and *Dacrydium* in Australia (Keppel et al., 2011). However, without knowing the phylogenetic relationship of Micropterygioideae to the rest of Lepidoziaceae, it is impossible to test hypotheses regarding its origins.

In this study, we infer the phylogeny of Lepidoziaceae using a multilocus data set that combines publicly available and newly generated molecular data. We perform a molecular dating analysis to infer the evolutionary timescale of the family, then use the dated tree to reconstruct its biogeographic history. Our study reveals the position of Micropterygioideae in the phylogeny of the Lepidoziaceae and provides an estimate of the divergence times and biogeographic history of the family.

## Materials and methods

### Molecular data set

To expand the taxonomic sampling of species of Lepidoziaceae, we sampled six herbarium specimens representing different species of *Micropterygium* from the Australian National Herbarium (CANB) (Supplementary Table S1). About 25 mg of dried tissue was cleaned from each sample and, from these tissues, DNA was extracted by the Australian Genome Research Facility (Brisbane). Following PCR amplification and amplicon purification, dual-direction Sanger sequencing of five markers by DNA BDT labelling reaction and capillary separation was carried out on an Applied Biosystems 3730xl Genetic Analyzer (Supplementary Table S2). We excluded sequences that did not show close affinity with available sequences from Lepidoziaceae, as assessed using BLASTn searches. We combined the resulting 14 sequences from *Micropterygium* with sequence data from 141 species of Lepidoziaceae and 30 outgroup taxa available on GenBank, to produce a data set comprising a total of 177 taxa. We included the outgroup taxa *Herbertus* and *Lepicolea*, which have been shown to be close relatives of Lepidoziaceae (Cooper et al., 2012b; Feldberg et al., 2014). We also included outgroup taxa representing other members of Jungermanniales (*Plagiochila, Calypogeia*, and *Scapania*) and Porellales (*Frullania, Acrolejeunea, Drepanolejeunea, Gackstroemia, Porella*, and *Radula*) to provide nodes for fossil calibrations.

Our assembled data supermatrix included nucleotide sequences from seven chloroplast markers, four mitochondrial markers, and two nuclear markers (Supplementary Table S3). This supermatrix had an occupancy of 44%, with 1013 sequences available out of a possible 2301 (13 markers from 177 taxa). We aligned the sequences of the 13 markers individually using MUSCLE (Edgar, 2004) and removed poorly aligned regions using Gblocks with less stringent selection (Castresana, 2000). We then tested for substitutional saturation and model adequacy using PhyloMad (Duchêne et al., 2018, 2022). Based on entropy scores calculated using only the variable sites, we removed three sequence alignments that were found to carry a high risk of misleading phylogenetic inference (first codon sites of *psbA*, first codon sites of *rbcL*, and *trnK*–*psbA* intergenic spacer).

### Phylogenetic analyses and molecular dating

We performed a phylogenetic analysis using maximum likelihood in IQ-TREE 2 (Bui et al., 2020), with the best-fitting partitioning scheme selected using a greedy search (Supplementary Table S4). Data subsets were allowed to evolve at different relative rates, such that the branch lengths were proportionate across subsets (Duchêne et al., 2020). Node support values were estimated using 1000 bootstrap replicates.

Using Bayesian phylogenetic analysis, we jointly estimated the phylogeny and divergence times in BEAST v2.7.3 (Bouckaert et al., 2019) using a birth-death tree prior and an uncorrelated lognormal relaxed clock (Drummond et al., 2006). Each data subset was allowed to have its own relative substitution rate. To calibrate the molecular clock, we specified a secondary calibration based on a previous age estimate for the split between Porellales and Jungermanniales (Laenen et al., 2014). We used a normal calibration prior with a mean of 319 Myr and standard deviation of 32.65 Myr (Ho and Phillips, 2009). In addition, we constrained the age of crown *Bazzania* to 34–381 Myr based on *Bazzania polyodus* (Feldberg et al., 2021), the only available fossil representing the family, and utilized 10 other fossil calibrations in the outgroup (Supplementary Table S5). All of these fossils can be unambiguously assigned to extant genera, and many have previously been used for setting a minimum age constraint on crown groups of the genera and subgenera of the two orders (Heinrichs et al., 2007; Cooper et al., 2012b; Feldberg et al., 2014). The maximum age constraint chosen for all fossil calibrations is based on a previous date estimate for the Porellales–Jungermanniales split (Laenen et al., 2014).

We partitioned the alignments according to the scheme selected in IQ-TREE and used Bayesian model averaging for all data subsets (Bouckaert and Drummond, 2017). The posterior distribution was estimated using Markov chain Monte Carlo sampling, with samples logged every 10^4^ steps over a total of 10^8^ steps. We ran the analysis three times and checked for sufficient sampling and convergence among the three chains using Tracer 1.7.1 (Rambaut et al., 2018). To examine any potential interactions among the calibration priors, we ran an additional analysis in which we sampled from the prior distribution.

### Biogeographical analyses

To investigate the historical biogeography of Lepidoziaceae, we obtained the distribution data of the species in the data set from authoritative literature sources, including taxonomic revisions and national flora treatments (Supplementary Table S6). We assigned the taxa to five floristic regions (Figure 2) based on Cox (2001): Holarctic, African, Indo-Pacific, South American, and Australian. The ancestral locations were inferred using Statistical Dispersal-Vicariance Analysis (S-DIVA) (Yu et al., 2010) in the software package RASP (Yu et al., 2011), allowing a maximum number of five areas at each node, because the long evolutionary history of the family spans the movement of the continents. To account for phylogenetic uncertainty, we performed the ancestral state reconstruction on the posterior sample of trees from our Bayesian analysis.

**Fig. 2.**
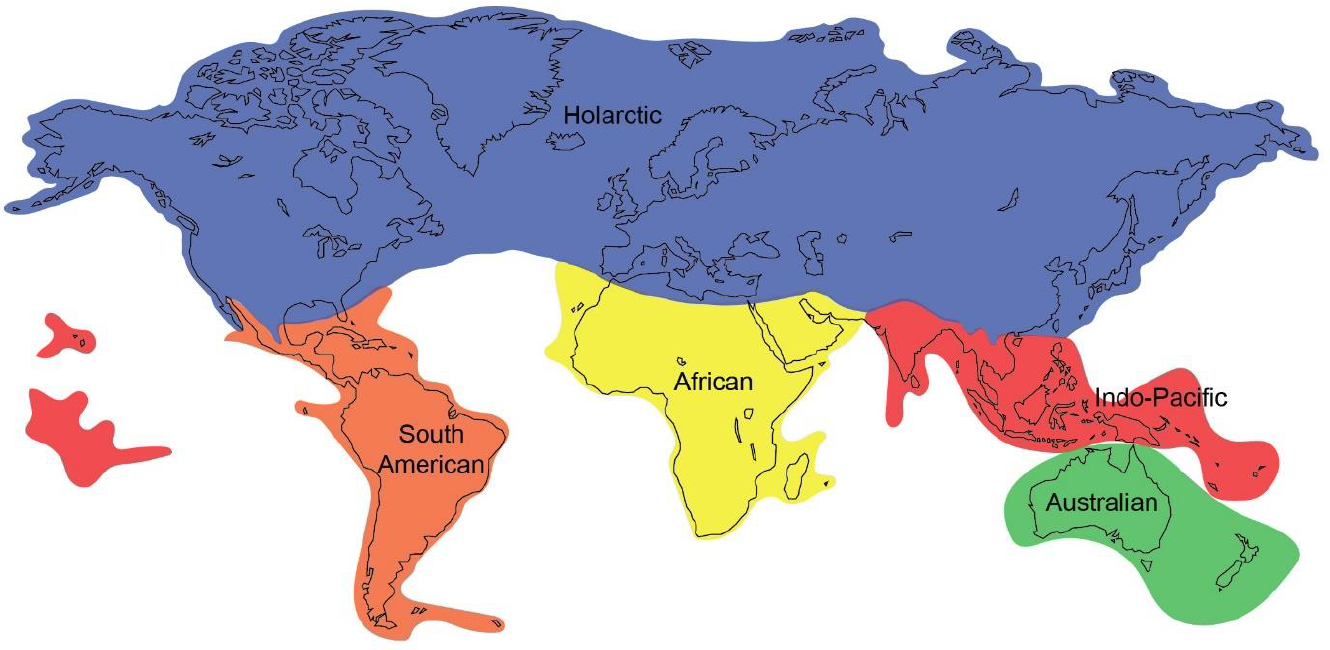
The five floristic regions defined by Cox (2001): African, Australian, Holarctic, Indo-Pacific, and South American. This definition of floristic kingdoms takes distinctiveness of flora (based on modern records) into account. The Australian Kingdom includes New Zealand in addition to Australia. Many representatives of Lepidoziaceae included in this study are found only in this kingdom. The Indo-Pacific and African Kingdoms are recognized here as separate kingdoms, in contrast to the floristic regions defined by Takhtajan (1986) who included both in the Paleotropical Kingdom. The South American Kingdom extends to the South Subantarctic islands. The Holarctic Kingdom, which remains unchanged from Takhtajan’s, includes North America, Eurasia, and North Africa.

## Results

### Phylogeny and divergence times

Our phylogenetic analyses yielded well-resolved trees for Lepidoziaceae with high bootstrap support and posterior probabilities for most nodes (Figure 3; Figure 4). Maximum-likelihood and Bayesian analyses supported the monophyly of the family. The six species of *Micropterygium* form a sister clade to Lembidioideae. Overall, the maximum-likelihood and Bayesian trees are similar to that inferred by Cooper et al. (2011) and support the same major clades, including Zoopsids I, Zoopsids II, and Zoopsids III of the polyphyletic subfamily Zoopsidoideae.

**Fig. 3.**
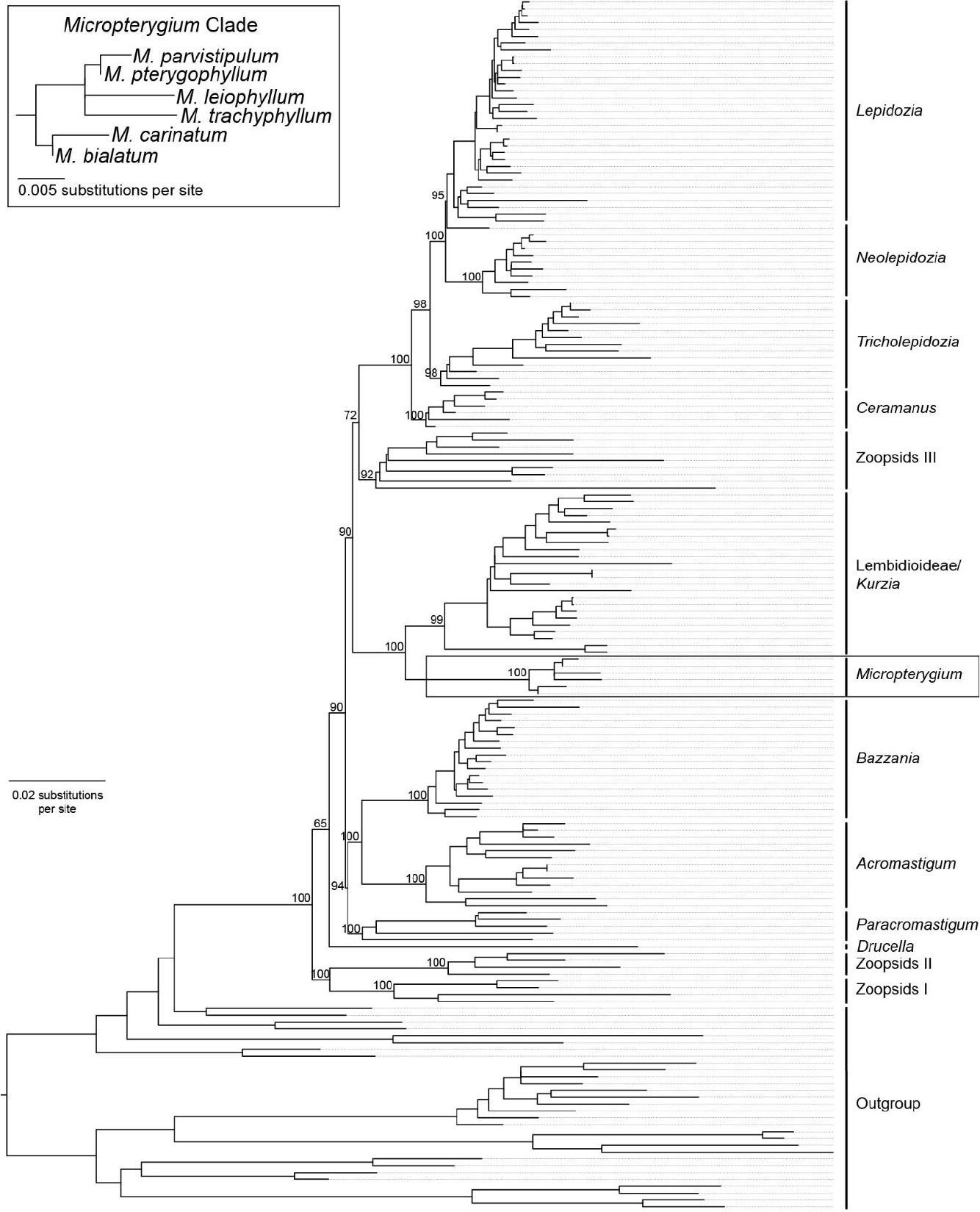
Maximum-likelihood tree of Lepidoziaceae inferred using a multilocus data set. The major branches are labelled with bootstrap support values. The inset shows a detailed view of the *Micropterygium* clade.

**Fig. 4.**
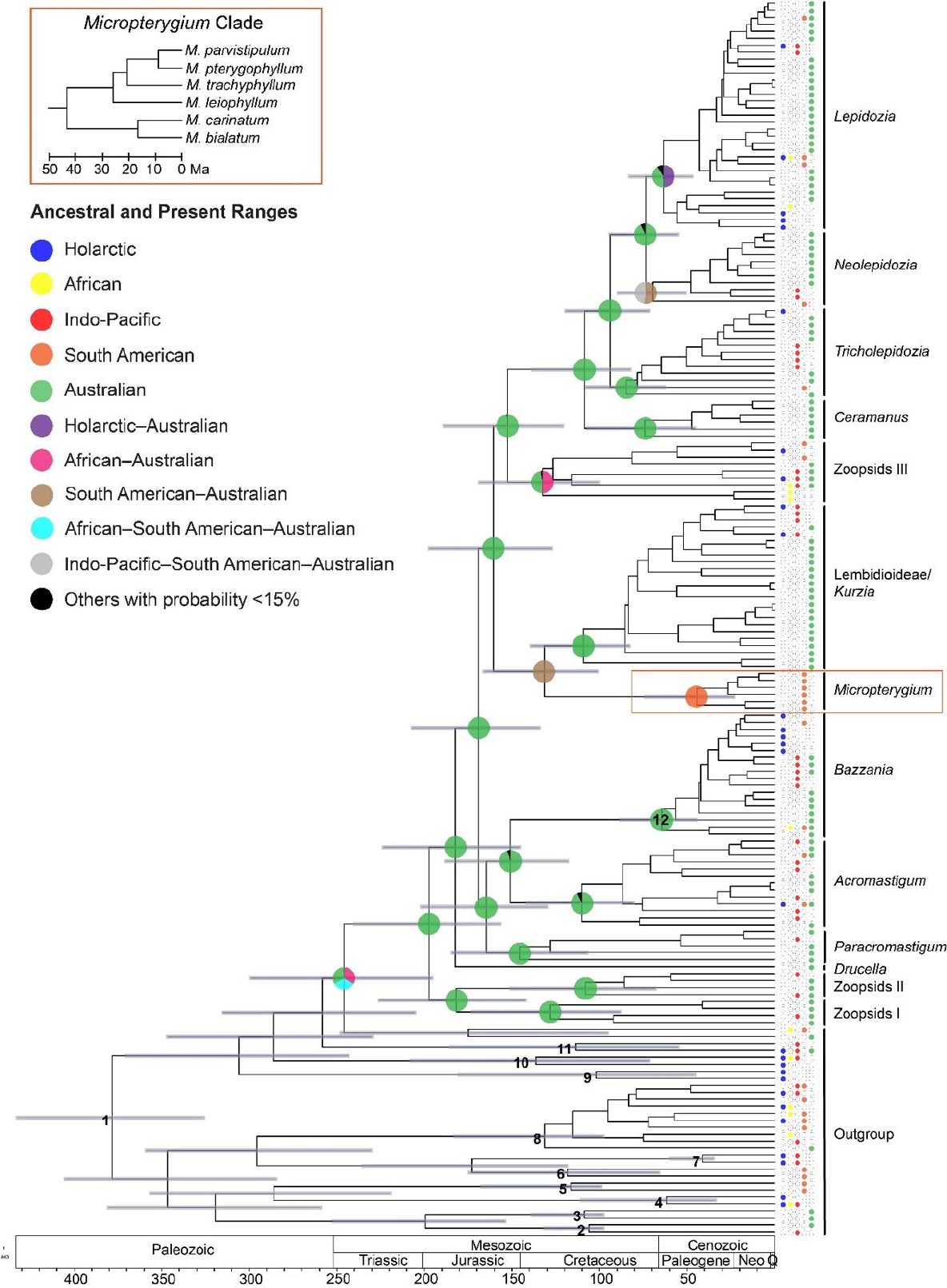
Dated phylogenetic tree of Lepidoziaceae inferred using a Bayesian relaxed-clock analysis of a multilocus data set. Circles at selected internal nodes show reconstructed ancestral ranges. Circles at the tips of the tree indicate present-day distributions. The inset shows a detailed view of the *Micropterygium* clade. Grey bars represent 95% credible intervals for the estimates of node ages. Numbers at internal nodes indicate the placement of calibrations for molecular dating: 1, secondary calibration based on Laenen et al. (2014); 2, *Radula* subg. *Amentuloradula* fossil; 3, *Radula* subg. *Odontoradula* fossil; 4, *Porella* fossil; 5, *Gackstroemia* fossil; 6, *Drepanolejeunea* fossil; 7, *Acrolejeunea* fossil; 8, *Frullania* fossil; 9, *Scapania* fossil; 10, *Calypogeia* fossil; 11, *Plagiochila* fossil; 12, *Bazzania* fossil.

Our results support the currently accepted circumscription of Lepidoziaceae (Schuster, 2000) as well as the revised circumscription presented by Cooper et al. (2012a) and Cooper (2013), where *Kurzia, Psiloclada*, and some species of *Telaranea* are excluded from Lepidozioideae. Moreover, Zoopsidoideae do not form a monophyletic group, with its genera appearing in separate clades (Zoopsids I, Zoopsids II, Zoopsids III, *Neogrollea*, and *Paracromastigum*). Bazzanioideae formed a monophyletic group with two distinct clades, *Acromastigum* and *Bazzania*. As in previous studies, some genera (including *Zoopsis, Telaranea*, and *Lembidium*) were found to be paraphyletic or polyphyletic.

The molecular dating analysis inferred a crown age for Lepidoziaceae of 197.3 Ma, with a 95% credible interval (CI) of 156.7–239.7 Ma (Figure 4). The split between Zoopsids I and Zoopsids II occurred 181.7 Ma (95% CI 142.2–225.7), whereas the divergence between Bazzanioideae and the *Paracromastigum* clade occurred 164.6 Ma (95% CI 130.0–201.7). The divergence between *Acromastigum* and *Bazzania* occurred 150.9 Ma (95% CI 118.2– 188.0). More recently, *Lepidozia* and *Neolepidozia* diverged from each other 73.6 Ma (95% CI 55.2–94.0), and the split between *Tricholepidozia* and the clade containing *Lepidozia* and *Neolepidozia* occurred 93.2 Ma (95% CI 71.6–119.2).

### Biogeographic reconstruction

Our reconstruction of ancestral location states placed the crown node of Lepidoziaceae in the Australian region and the divergence of the family from its sister lineage in three possible locations (Australian, African–Australian, and African–South American–Australian) (Figure 4). Most of the major clades also originated in the Australian region. Crown Micropterygioideae emerged in South America. The analysis yielded more than one possible location for the origins of crown *Lepidozia* (Australian and Holarctic–Australian), crown *Neolepidozia* (South American–Australian and Indo-Pacific–South American–Australian), and crown Zoopsids III (African–Australian and Australian).

## Discussion

### Phylogeny and age of Lepidoziaceae

Our phylogenetic analysis of a multilocus data set has resolved the evolutionary relationships among major lineages within the family Lepidoziaceae, including the placement of the subfamily Micropterygioideae. The maximum-likelihood and Bayesian trees inferred in our study are congruent with that of the most recent molecular phylogenetic study that used a large data set (Cooper et al., 2011), where the monophyly of the family was strongly supported. The same separate clades of Zoopsidoideae were inferred here, confirming the polyphyly of that subfamily.

Our molecular dating analysis, based on 11 fossil calibrations and one secondary calibration, yielded an age estimate for crown Lepidoziaceae of 197.3 Ma (156.7–239.7). Our secondary calibration was based on a date estimate by Laenen et al. (2014), which we chose over other possible sources for secondary calibration because it represents the most comprehensive integration of fossil data (35 moss, 25 liverwort, and three hornwort fossils) and molecular data (eight markers: five chloroplast, two mitochondrial, and one nuclear) among all previous studies that estimated the divergence times among major liverwort groups. The posterior date estimate of the Porellales–Jungermanniales split, at 378.4 (326.2– 432.5) Ma, is much older than that specified in the calibration prior (mean=319; standard deviation=32.65). This shift appears to be driven by a signal in the molecular data and the inclusion of a range of fossil-based minimum age constraints across the tree but is not the product of interactions among the calibration priors (Supplementary Figure S1).

Our inferred divergence times for the family Lepidoziaceae are also much older than previous estimates. The analysis by Feldberg et al. (2014), which estimated the crown age of Lepidoziaceae at 174 Ma, used 20 fossil calibrations and a data set of 303 liverwort species (22 Lepidoziaceae). The phylogenetic tree was inferred using an unpartitioned analysis of *rbcL*. The study by Cooper et al. (2012b), which estimated the crown age of Lepidoziaceae at 116 Ma, used nine fossil calibrations and a data set comprising only three molecular markers from 212 liverwort species (64 Lepidoziaceae). Although that study used a partitioned data set, the absolute ages of the fossils were not used in the calibrations.

### Evolutionary timescale and biogeographic history

The results of our molecular dating and biogeographic analyses allow us to propose an account of the evolutionary history of Lepidoziaceae. The crown age of the family is estimated at 197.3 Ma (156.7–239.7), after the break-up of Pangaea but before the early fragmentation of Gondwana. Furthermore, many of the major lineages, including Bazzanioideae, Lepidozioideae, Lembidioideae, Zoopsids I, Zoopsids II, Zoopsids III, and *Paracromastigum*, have estimated crown origins before Africa split from Antarctica during the mid-Cretaceous (McLoughlin, 2001), leaving South America, Australia, and New Zealand still connected to Antarctica (Smellie et al., 2020; van den Ende et al., 2017). These estimated ages suggest that these lineages of the family had sufficient time to spread throughout Gondwana before it began to break up. The results of our biogeographical analyses support an origin in land masses that were part of Gondwana, as postulated by Schuster (2000), considering that the inferred ancestral range of crown Lepidoziaceae is Australian and that the inferred possible locations of the divergence of the family from its sister lineage all show continents previously part of Gondwana (Figure 4). Furthermore, many lineages (Zoopsids I, Zoopsids II, *Paracromastigum*, Bazzanioideae, Lembidioideae/*Kurzia*, and Lepidozioideae) have inferred crown origins in the Australian region.

Although vicariance events could potentially account for the key divergences in Lepidoziaceae, the same cannot be said for some of the species-level divergences. For instance, the lineage leading to *Z. argentea*, which occurs on Sunda in addition to Australia (part of Sahul) and New Zealand, may have diverged from *Z. nitida* about 92.1 Ma (52.8–134.1 Ma), much earlier than the formation of Sunda islands in the Miocene (Hall, 2002), and must have reached Sunda from Sahul by means of dispersal. The conditions in the mid-Miocene until present might have favoured floristic exchange between the two shelves through dispersal (Crayn et al., 2015). Although the estimated divergence of *Lepidozia* from *Neolepidozia* about 73.6 Ma (55.2–94.0) predates the separation between Australia and New Zealand (Veevers and McElhinny, 1976), the occurrence of *Lepidozia* in all five floristic regions is incompatible with vicariance. Direction-dependent long-distance dispersal by wind has been found to be responsible for floristic similarities in the Southern Hemisphere (Muñoz et al., 2004) where nearly all genera of Lepidoziaceae are present. Biological dispersers such as birds (Proctor, 1961; Fife and de Lange, 2009; Chmielewski and Eppley, 2019) and bats (Parsons et al., 2007) might also have had a strong influence on past and present distribution patterns.

### Micropterygioideae and its circum-Antarctic links to Lepidoziaceae

Our analysis resolved the neotropical endemic subfamily Micropterygioideae as the sister group to Lembidioideae and supports their status as a separate subfamily. A close relationship between Micropterygioideae and Lembidioideae is supported by morphological similarities mentioned by Schuster (2000). *Lembidium nutans* has loosely folded leaves resembling half canoes, similar to those of *Micropterygium*. The oil bodies in the species of both subfamilies are either reduced or completely lacking. *Micropterygium* was divided into two subgenera (Schuster, 2000), namely, subg. *Pseudolembidium* and subg. *Micropterygium*, without division into sections. All six species of *Micropterygium* in this study are in the latter subgenus, which is characterised by anisophyllous leaves. In another subgeneric classification scheme, the genus was divided into three sections (Reimers, 1933), namely, sect. *Conchifolia*, sect. *Subaequifolia*, and sect. *Genuina*. From the species included in the data set, *M. leiophyllum, M. parvistipulum, M. pterygophyllum*, and *M. trachyphyllum* are all included in sect. *Genuina*, whereas the other two sections are unrepresented. *Micropterygium bialatum* and *M. carinatum*, which form a sister group to the rest of the genus, are not included in any of these sections. A more comprehensive sampling of the genus is needed to test the subgeneric classification schemes that have been proposed. The phylogenetic position of the monotypic genus *Mytilopsis* remains to be resolved. Further sampling of Lepidoziaceae will allow resolution of the remaining phylogenetic uncertainties.

The restricted range of Micropterygioideae in the Neotropical region stands in contrast with the wide distribution of the family as a whole, and this could, perhaps, be explained by the factors that limit dispersal, establishment of spores, and population growth. Liverworts show no correlation between spore size and range (Laenen et al., 2016), but a strong correlation has been found between range and asexual reproduction. The lack of asexual reproduction in the subfamily (Schuster, 2000), as well as its relatively young crown age, also provide potential explanations for its limited geographic distribution. It is also possible that the lineage once occupied the Australian region and then went extinct there at some point. The results of our phylogenetic dating analysis support this possibility. We estimated that Micropterygioideae split from Lembidioideae 131.6 Ma (101.1–165.8) in the Australian and South American regions during a time when Antarctica was still connected to Africa, South America, and Australia. The estimated crown origin of the subfamily is 44.6 Ma (23.3–73.9) in South America, suggesting that it has occurred before South America separated from Antarctica 30 Ma (van den Ende et al., 2017). These circum-Antarctic links of the subfamily to the rest of the family strongly suggests extinction of the lineage in the region, but this requires verification through fossil evidence.

Neotropical endemic taxa either have origins in the Neotropics itself, e.g., the angiosperm family Calyceraceae (Brignone et al., 2023), or from elsewhere, e.g., the fern family Cyatheaceae (Korall and Pryer, 2014). Our study shows that Micropterygioideae are among the lineages that the Neotropical region holds in its collection of taxa with circum-Antarctic links. There are several other Lepidoziaceae taxa that are exclusive to the Neotropics, including *Protocephalozia* of the monotypic Protocephalozioideae and some elements of the heterogeneous Zoopsidoideae (*Monodactylopsis, Odontoseries*, and *Pteropsiella*). Including these taxa in the phylogenetic dating of Lepidoziaceae will reveal their biogeographical connection to the rest of the family.

## Conclusions

Our study has presented a reconstruction of the evolutionary and biogeographic history of the liverwort family Lepidoziaceae, including the phylogenetic placement of the subfamily Micropterygioideae. Key divergences can be explained by vicariance, but long-distance dispersal is likely to have played a large role in the recent diversification of the family. However, the phylogeny of Lepidoziaceae is still not fully resolved. Improved resolution of phylogenetic relationships and biogeographic history can potentially be achieved through more comprehensive taxon sampling, with genetic data yet to be obtained from several genera in the family. To allow confident taxonomic revision, it will be necessary to conduct further phylogenetic analysis using larger numbers of markers, such as those obtained by exon capture, transcriptomics, or even whole-genome sequencing.

## Acknowledgements

We thank Brendan Lepschi, Christine Cargill, Endymion Cooper, and Judith Curnow of the Australian National Herbarium (Canberra) for their kind assistance during the acquisition of the samples used in this study. A.L.R. was supported by a Postgraduate Student Travel Grant administered by the Australasian Systematic Botany Society Inc. (supported by the Australian Biological Resources Study) and by the Foreign Graduate Scholarship Program from the Science Education Institute of the Department of Science and Technology (DOST-SEI) of the Philippines.

## Notes

### Competing Interest Statement

The authors have declared no competing interest.

