## Supplementary Material for "The Neotropical endemic liverwort subfamily Micropterygioideae had circum-Antarctic links to the rest of the Lepidoziaceae during the early Cretaceous"

**Table S1.** *Micropterygium* samples from the Australian National Herbarium (CANB) used in this study.

| Species | Collection number | Accession number |
|---|---|---|
| *M. bialatum* | Boom and Gopaul 7447 | CANB 00906825 |
| *M. carinatum* | Halling 5534 | CBG 9210343 |
| *M. leiophyllum* | Prance et al. 15566 | CBG 8205487 |
| *M. parvistipulum* | Prance et al. 11404 | CBG 8205491 |
| *M. pterygophyllum* | Buck 11370 | CANB 00785805 |
| *M. trachyphyllum* | Boom and Gopaul 7240 | CANB 00906826 |

**Table S2.** Molecular markers amplified and sequenced for the *Micropterygium* samples in this study.

| Marker | Genome | Primer Sequences (5′-3′) | Reference |
|---|---|---|---|
| trnL-trnF | chloroplast | F: ATTTGAACTGGTGACACGAG<br>R: CGAAATCGGTAGACGCTACG | Taberlet et al. (1991) |
| trnG intron | chloroplast<br>chloroplast | F: ACCCGCATCGTTAGCTTG<br>or ATTCGGTGATTTAGTTACG*<br>R: GCGGGTATAGTTTAGTGG | Pacak and Szweykowska-Kuliñska (2000) |
| psbA-trnH | chloroplast | F: GTTATGCATGAACGTAATGCTC<br>R: CGCGCATGGTGGATTCACAATCC | Stech et al. (2011) |
| nad1 | mitochondrial | F: GCATTACGATCTGCAGCTCA<br>R: GGAGCTCGATTAGTTTCTGC | Sun (2002) |
| 26S rDNA | nuclear | F: GAGTCGGGTTGTTTGGGA<br>R: TTGGTCCGTGTTTCAAGACG | Kuzoff et al. (1998) |

*Forward primer redesigned to target shorter amplicon for higher chance of success

**Table S3.** Data supermatrix (Microsoft Excel file provided).
https://docs.google.com/spreadsheets/d/1MWUVFd9Dk3wNh90I5XJPiEeQtORnlZwXCKOsh2qU1Ow/edit#gid=0

**Table S4.** Partitioning scheme used for maximum-likelihood and Bayesian phylogenetic analyses.

| **Subset** | **Substitution model** | **Markers** |
|---|---|---|
| 1 | TN+F+I+G4 | 26S rRNA, atpB codon position 1 |
| 2 | GTR+F+I+G4 | 5.8S rRNA, nad1 |
| 3 | GTR+F+G4 | ITS1, ITS2 |
| 4 | GTR+F+I+G4 | atpB codon position 2, psbA codon position 2, rbcL codon position 2 |
| 5 | TVM+F+I+G4 | atpB codon position 3, rbcL codon position 3, rps4 codon position 3 |
| 6 | GTR+F+I+G4 | nad5-nad4, nad5, rps4 codon position 1, rps4 codon position 2 |
| 7 | TIM+F+I+G4 | psbA-trnH intergenic spacer, trnG intron |
| 8 | GTR+F+I+G4 | psbA codon position 3 |
| 9 | K3Pu+F+I+G4 | psbT-psbH, trnL-trnF |
| 10 | GTR+F+I+G4 | rps3 |

**Table S5.** Fossils used for age calibrations based on Feldberg et al. (2014).

| Fossil | Age assignment | Corresponding taxon | Origin and age |
|---|---|---|---|
| *Drepanolejeunea eogeana* | 15–381 | *Drepanolejeunea* crown | La Toca Formation, Dominican Republic; 15–20 Ma |
| *Calypogeia stenzeliana* | 34–381 | *Calypogeia* crown | Baltic region; 34–41 Ma |
| *Porella subgrandiloba* | 34–381 | *Porella* crown | Baltic region; 34–41 Ma |
| *Bazzania polyodus* | 34–381 | *Bazzania* crown | Baltic region; 34–41 Ma |
| *Plagiochila groehnii* | 34–381 | *Plagiochila* crown | Baltic region; 34–41 Ma |
| *Scapania hoffeinsiana* | 34–381 | *Scapania* crown | Baltic region; 34–41 Ma |
| *Acrolejeunea ucrainica* | 35–381 | *Acrolejeunea* crown | Klesov, Ukraine; 35–37 Ma |
| *Gackstroemia cretacea* | 99–381 | *Gackstroemia* crown | Northern Myanmar; 99 Ma |
| *Frullania baerlocheri, F. cretacea, and F. partita* | 99–381 | *Frullania* crown | Northern Myanmar; 99 Ma |
| *Radula cretacea* | 99–381 | *Radula* subg. *Odontoradula* crown | Northern Myanmar; 99 Ma |
| *Radula heinrichsii* | 99–381 | *Radula* subg. *Amentuloradula* crown | Northern Myanmar; 99 Ma |

**Table S6.** Distribution data (Microsoft Excel file provided).
https://docs.google.com/spreadsheets/d/1FjlGvI-M8awSS6WQ4ZeuatAqzDcQN6UJwJDuaDhKE8Y/edit#gid=1219347240

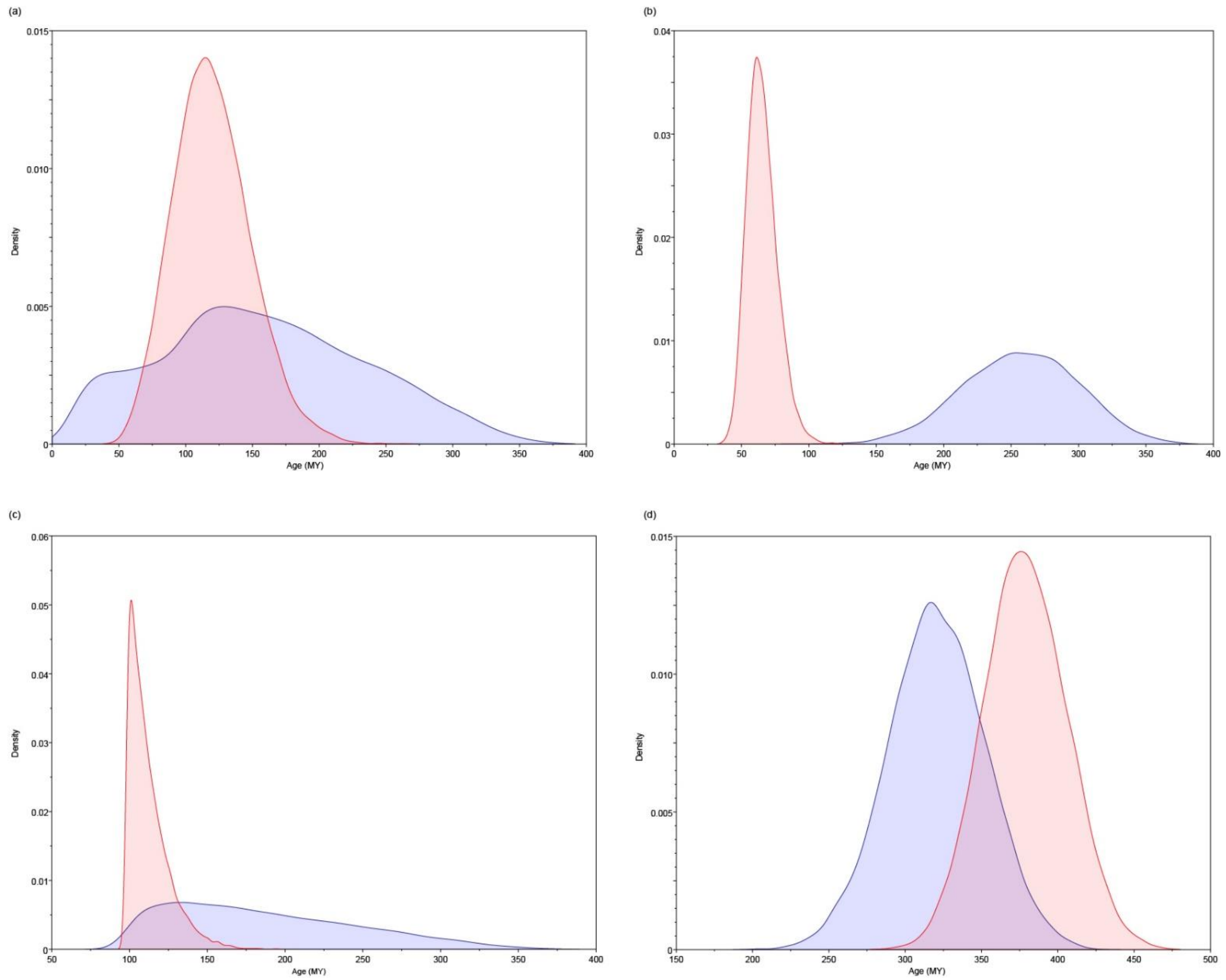


**Figure S1.** Comparisons of marginal prior densities (blue) and marginal posterior densities (red) for the ages of four nodes in the phylogenetic tree. (a) *Drepanolejeunea* crown, (b) *Bazzania* crown, (c) *Radula* subg. *Odontoradula* crown, and (d) Porellales–Jungermanniales split.
